# Mapping corpus callosum architecture: developmental, genetic, and cognitive correlates in youth

**DOI:** 10.64898/2026.04.25.720714

**Authors:** Vanessa Siffredi, Léa Schmidt, Robin J. Hofmeister, Jonathan Patino Lopez, Zoltán Kutalik, Jonas Richiardi

## Abstract

The corpus callosum (CC), the largest interhemispheric white-matter tract, plays a central role in higher-order cognitive functions and is frequently altered in neurodevelopmental and psychiatric conditions. However, most diffusion MRI studies rely on tract-averaged measures, which obscure spatially specific microstructural variation along the tract that may hold biologically and functionally meaningful information. This study aimed to provide a fine-grained characterisation of spatial variation in callosal structure and to determine its developmental, genetic, and cognitive correlates.

This study leveraged multimodal data from the Philadelphia Neurodevelopmental Cohort (PNC), comprising 1342 participants aged 8–21 years. Diffusion MRI and tractography-based segmentation were used to extract fine-grained spatial variation in diffusion metrics along seven callosal tracts. A candidate gene approach targeted four genes (*DCC*, *CDH2*, *AKT3*, and *GLI3*) previously linked to the neurobiological mechanisms underlying CC formation. Five cognitive factors were derived from the Penn Computerized Neurocognitive Battery using factor analysis. A two-stage functional data analysis approach to examine global and local associations between along-tract diffusion metrics and age, candidate genetic variants, and behavioural outcomes.

Results revealed distinct midline-to-cortical variations of age-related diffusion metrics change across callosal subdivisions. Frontal and parietal heteromodal callosal pathways showed pronounced distal-segment maturation (F = 13 – 23, p ≤ 2.8×10e-16), whereas posterior sensorimotor and occipital callosal tracts exhibited more stable age associations along their lengths (F = 4.2 – 4.3, p = 10e-3). Genetic variations in callosal axon-guidance genes (ROBO1, IQCJ-SCHIP1, NRP1 and DCC) were associated with spatial variation in callosal diffusion metrics, particularly in anterior (rostrum) and posterior callosal subdivisions (isthmus and splenium) (F = 4.29 – 18.61, p-values = 2.2e-16 – 1.4e-04). These regions correspond to early-forming callosal compartments, suggesting that prenatal axon-guidance mechanisms leave enduring spatial patterns on callosal organisation. Finally, spatial variation in callosal microstructure was significantly associated with behavioural performance (F = 2.9 – 21.3, *P* = 2.8×10e-16 – 0.04), with the strongest and most spatially heterogeneous effects observed for complex cognition and executive functioning. Across all analyses, functional data models generally outperformed tract-averaged linear models, supporting the value of explicitly preserving spatial variation along callosal tracts.

Our findings converge on the CC as a spatially differentiated structure in which early genetic modulators and developmental constraints shape region- and segment-specific microstructural architecture that is behaviourally relevant through childhood and adolescence.

## Introduction

With more than 250 million axons, the corpus callosum (CC) is the largest commissural bridge of white-matter bundles connecting the left and right hemispheres in the human brain. As the primary cortical projection system, structural alterations of the CC can exert widespread effects on brain organisation, influencing not only its own axonal architecture but also bilaterally connected cortical regions and broader patterns of structural asymmetry and functional lateralisation.^1–3^ Serving as an information highway, the CC supports the integration and coordination of both low-level sensory-motor processes and higher-order cognitive functions across the two hemispheres. Consistent with the centrality of this brain structure, variation in CC architecture has been associated with key cognitive functions, ranging from executive and attentional functions to memory, language, and socio-emotional abilities.^4–6^ Moreover, alteration in CC architecture have been consistently reported in neurodevelopmental and neuropsychiatric disorders, including autism spectrum disorder,^7–9^ attention-deficit/hyperactivity disorder ^10,11^ and schizophrenia. ^12,13^

The formation of the CC is governed by complex neurobiological mechanisms that unfold through a highly coordinated sequence of developmental events.^14^ During early embryogenesis, all telencephalic commissures initially cross the interhemispheric midline through the commissural plate.^15^ At approximately 11–12 weeks postconception, pioneering axons arising from the cingulate cortex extend along a guided path and are the first to cross this midline plate.^15–17^ After crossing, these axons continue into the contralateral hemisphere toward their target regions and provide guidance cues for later callosal fibres emerging from the neocortex.^6,18^ During this axonal guidance phase, several molecular signalling pathways are crucial for directing callosal fibre growth. Among them, NRP1 (neuropilin-1) encodes a receptor that mediates growth cone responses to guidance cues,^19–21^ while DCC encodes the DCC receptor that interacts with netrin-1 to regulate midline guidance.^22–24^ In parallel, ROBO1, a key receptor in the SLIT–ROBO signalling pathway, provides repulsive cues that guide callosal axons toward the midline while preventing entry into inappropriate ventral regions.^25^ Disruption of ROBO1 signalling in mouse models leads to severe callosal misrouting and midline crossing defects.^26,27^ Complementing these guidance receptor systems, emerging evidence highlights the role of the intracellular structural regulator for axonal guidance. IQCJ-SCHIP1, a brain-enriched isoform of the SCHIP1 gene, contributes to axonal organisation by stabilising molecular complexes at the axon initial segment and nodes of Ranvier, supporting calcium-dependent signalling, and regulating axon outgrowth and responsiveness to guidance cues.^28,29^ Recent large-scale genetic analyses identified IQCJ- SCHIP1 as a key contributor to corpus callosum morphology,^30^ and loss-of-function studies in animal models show that SCHIP1 disruption leads to abnormal axon trajectories and defects in commissural pathway formation.^31^ Although these molecular mechanisms operate during a relatively restricted prenatal window, they lay the foundation for subsequent callosal development. By around 20 gestational weeks, the basic structural layout of the corpus callosum is established,^14,15,32^ but refinement of callosal connections and myelinisation continue throughout the third trimester and extend well into childhood and adolescence.^4,33^ Perturbations in axonal guidance during early development therefore have cascading downstream effects on the structural organisation of callosal white-matter tracts observed later in life.^20,21,34–36^ Consistent with this, variants in these callosal guidance-related genes have been associated with increased vulnerability to neurodevelopmental and psychiatric disorders, including autism spectrum disorder ^37–40^ and attentional deficit hyperactivity disorder.^41,42^

Diffusion magnetic resonance imaging (dMRI) offers a unique opportunity to investigate white-matter characteristics non-invasively in vivo.^43^ This approach has enabled large-scale studies to map changes in white-matter characteristics throughout childhood and adolescence. Most commonly, dMRI studies estimate white-matter microstructure using metrics derived from diffusion tensor imaging (DTI), which models the diffusion of water in tissue. Despite its limitations,^44^ DTI has been widely used because it yields quantitative measures, including fractional anisotropy (FA) and mean diffusivity (MD), recognised to reflect key aspects of white-matter microstructure, including myelin thickness, local fibre density, fibre packing and organisation, and overall tissue “integrity”.^45,46^ These measures have also been found to be sensitive to developmental changes,^47–50^ modulated by genetic variation ^51^ and associated with individual differences in cognitive and socio-emotional functioning.^10,52–54^ Nonetheless, most developmental dMRI studies rely on tract-averaged diffusion metrics, reporting global age-related increases in FA during childhood or tract-level group differences in paediatric populations, such as preterm-born individuals or individuals with ADHD compared to typically developing peers.^48–50,55–59^ Such averaging approaches at the tract-level assume uniform microstructural organisation along the tract. In contrast, evidence from post-mortem, animal, and recent neuroimaging studies indicates substantial along tracts variation in white-matter maturation.^50,57,58^ In the corpus callosum specifically, heterogeneity in early axon guidance and cortical projections may give rise to marked along-tract differences in microstructural characteristics.^59^ Accounting for such along-tract spatial heterogeneity may therefore provide a more biologically precise understanding of how developmental, genetic, and cognitive factors relate to callosal microstructural organisation.

To move beyond tract-averaged approaches and building on evidence that spatial heterogeneity along the corpus callosum holds biologically and functionally meaningful information, this study aimed to provide a fine-grained characterisation of spatial variation in callosal structure and to determine its developmental, genetic, and cognitive correlates. More specifically, leveraging multimodal data from the Philadelphia Neurodevelopmental Cohort (PNC) (ages 8–21) and a functional data analysis framework, we characterised the developmental trajectory of spatial variation in callosal microstructure, examined the contribution of early callosal axon guidance genes to these spatial patterns, and assessed their associations with behavioural outcomes.

## Materials and methods

### Participants

The Philadelphia Neurodevelopmental Cohort (PNC) is a publicly available population-based sample, with participants between 8 and 21 years from the greater Philadelphia area.^60,61^ It includes clinical, genetic and cognitive data, as well as neuroimaging for a subset. Recruitment procedures, sample characteristics, clinical, cognitive, and imaging protocols are described in detail by Satterthwaite et al (2016). The institutional review boards of the University of Pennsylvania and the Children’s Hospital of Philadelphia approved all study procedures, and written informed consent was obtained from all participants.

The diffusion MRI (dMRI) dataset consisted of 1342 participants. Of these participants, 294 participants were excluded due to excessive head motion during acquisition, defined as mean framewise displacement (FD) > 0.5 mm or maximum FD > 6 mm (details in section below); all preprocessing and reconstructions underwent visual quality control.^43,62,63^ Participants without both usable dMRI data and completed behavioural assessments (n = 5) were also excluded. The final sample comprised 1043 participants (566 females [54.3 %]) aged 8 to 21 years at scan (mean [SD] age = 15.74 [3.41] years).

### dMRI acquisition, processing and tract segmentation

dMRI scans were acquired at University of Pennsylvania on a 3-T Siemens TIM Trio scanner (Siemens Medical Solutions), acquired using a twice-refocused spin-echo single-shot echo-planar imaging sequence sequence (FoV 240 × 240 mm; matrix 128 × 128 × 70; 64 gradient directions; b = 1000 s/mm2; voxel size 1.875 × 1.875 × 2 mm).^60^

dMRI data preprocessing and reconstruction were performed using QSIPrep 0.19.1,^64^ which is based on Nipype 1.8.6 (^65,66^ RRID:SCR_002502), Figure 1. White matter bundles of the corpus callosum were segmented using TractSeg ^67^ for seven callosal subdivisions: Rostrum (denoted CC 1), Genu (CC 2), Rostral Body (CC 3), Anterior Midbody (CC 4), Posterior Midbody (CC 5) and Isthmus (CC 6) and the Splenium CC (CC 7). Detailed preprocessing, reconstruction and tract segmentation can be found in the supplementary material.

**Figure 1.**
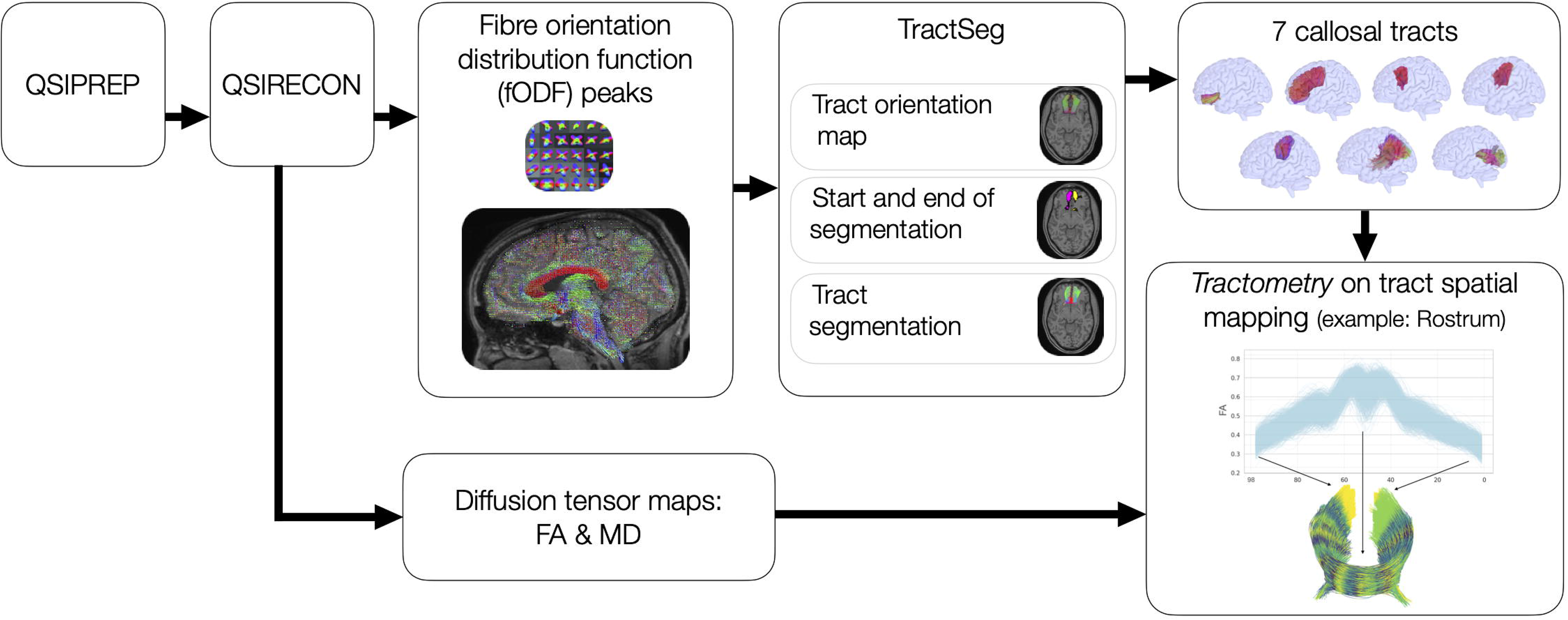
Diffusion image processing pipeline. Flowchart summarising diffusion image preprocessing, model fitting, tractography and tractometry measures. CC, corpus callosum; FA, Fractional Anisotropy; MD, Mean Diffusivity. In short, after preprocessing with QSIPrep, fibre orientation distribution functions (fODFs) were estimated using QSIRecon. These voxelwise fODF peaks were used by TractSeg to segment seven corpus callosum bundles through deep-learning–based identification of bundle orientation, start–end regions, and streamline reconstructions. In parallel, fractional anisotropy (FA) maps were computed from DTI fitting using QSIRecon. Using Tractometry, each callosal tract was sampled into an equal number of equidistant segments (*n=*98) and for each segment, the average FA of all streamlines was calculated, yielding a fine-grained curve of FA values along each tract. An example FA along-tract spatial variation is shown for the rostrum of the CC.

### Tractometry

Each CC white-matter tract was divided into 100 segments using the assignment mapping approach described by Chandio et al. (2020) ^68^ and implemented into TractSeg. Every point of the tract was assigned to the closest points of the centroid on the model tract to create assignment maps/segments. Assignments are created in a common space (model tract), which ensures that the segment index corresponds to the same centroid across all individuals. A model tract is represented with 100 centroids uniformly distributed along the length of each tract in a common reference space (for more detailed procedure, see ^68^). Importantly, this method uses all original points of the tracts, avoiding artificial resampling or reshaping. Tract segmentations were then applied to the diffusion metrics of interest, i.e., FA and MD maps, using the Tractometry implementation in TractSeg to segment-wise measures of callosal white-matter microstructure. This yielded 98 along-tract FA and MD values per tract (first and last segments are automatically removed due to unreliable estimates at tract endpoints).

### Genetic data processing and analysis

Genotype data were obtained from dbGaP (phs000607.v2.p2). Biological samples (i.e., blood sample) from PNC participants who also completed the MRI were genotyped by the Center for Applied Genomics at the Children’s Hospital of Philadelphia. Genotype were completed across participant in 8 batches using 7 different types of Affymetrix and Illumina arrays (i.e., Axiom, and Illumina HumanHap550 (v1, v3), Human1M-Duo (v3), Human610-Quad (v1), Genome-Wide Human SNP Array 6.0 and HumanOmniExpress).^69^ Supplementary fTable S2 provide sample and SNP counts by genotyping array before and after quality control, imputation and merge of genotype data. Detailed quality control, data cleaning, imputation, merge of genotype data, genetic ancestry and population structure correction, and candidate gene coordinates extraction are available in the Supplementary material. In short, genotype data were processed by array batch using PLINK v2.0 ^70^ to perform standard quality control, excluding individuals and SNPs with >5% missingness, low minor allele frequency <1%, or Hardy–Weinberg equilibrium p-value <1x10e-6. We harmonised genome build ^71^ and allele coding across datasets, aligned strands to the 1000 Genomes Phase 3 reference panel,^72^ normalised alleles, and performed haplotype phasing with SHAPEIT5 using genetic maps and chromosome-specific reference haplotypes from the 1000 Genomes Phase 3 panel (aligned to the GRCh37 human genome assembly).^73^ Phased genotypes were imputed separately by dataset and chromosome using IMPUTE5 with the 1000 Genomes Phase 3 reference panel and a b37 recombination map. High-quality imputed variants (imputation INFO Score > 0.8) were retained, and normalized VCF files were sorted, indexed, and merged across genotyping platforms into a single multi-array dataset using bcftools. An unrelated sample was constructed and used for all subsequent analyses: for each family, the participant with the highest genotyping completeness across the four candidate gene regions was retained. To account for population stratification,^74^ genetic ancestry was assessed using principal component analysis (PCA) on the merged imputed genotype dataset. Based on the examination of the variance explained by each PC (Supplementary Figure S2) and previous studies,^75^ the first ten ancestry PCs were included as covariates in subsequent statistical models. As described in the introduction section, four genes implicated in callosal axonal guidance were selected for the current candidate gene association study: NRP1, DCC, ROBO1 and IQCJ-SCHIP1. Gene coordinates (chromosome, start, end) were obtained from the Ensembl GTF annotation file within the gene body and extended by ±100 kb flanking regions.

Monomorphic SNPs and, where applicable, fewer than 100 complete genotype–phenotype observations were excluded. For each candidate gene, LD matrices were computed as Pearson correlation coefficients (r) between SNP pairs using R (v4.4.3) (Supplementary Figure S3). To correct for multiple testing while accounting for LD correlation among SNPs, we applied an LD-aware Bonferroni correction per gene. Rather than assuming independence across all tested variants, the effective number of independent tests (Meff) was derived from LD matrices using Gao’s method, implemented in the SimpleM R package, and used to compute corrected significance thresholds ^76–78^ (Supplementary Table S3). To identify independent association signals within each candidate gene region, significant variants (p value < candidate gene-specific LD-aware threshold) were subsequently clumped using snp_clumping from the bigsnpr R package,^79^ with in-sample LD computed from study genotype data, an r² threshold of 0.1, and a 100 kb window.

### Behavioural measures

Cognitive abilities were measured using the Penn Computerized Neurocognitive Battery (CNB), including 12 computerised tests spanning verbal and nonverbal reasoning, executive control and working memory, episodic memory, and social cognition.^80,81^ These included accuracy and reaction time measures. Factor analyses were completed following the procedure of previous studies using the same PNC data.^80,82^ For the 1043 participants with valid CNB scores and less than 50% missing data, missing values were imputed using multiple imputation by chained equations (MICE; R package mice). To align the direction of interpretation across measures, reaction time scores were inverted, such that higher scores indicated better performance. All cognitive scores were regressed on age at testing and sex yielding standardised residuals z-scores. For each of the 12 cognitive tasks, efficiency scores were computed by combining the accuracy and reaction-time z-scores (Efficiency = Accuracy_z + RT_z). This yielded a single composite index reflecting both response accuracy and processing speed. A factor analysis was performed on the efficiency scores guided by previous findings supporting a four-factor structure.^80^ The four neurocognitive factors correspond to four domains: complex cognition, executive functioning, memory and social cognition (Supplementary Figure S1, Supplementary Table S1). Factor scores for each participant, derived using the loading matrix and standardised efficiency variables, were then used in subsequent analyses and referred as behavioural scores: complex cognition, executive functioning, memory and social cognition.

### Statistical analyses

#### Along-tract functional data analysis approach

Statistical analyses are described in detail in the Supplementary Material. In brief, we used a two-stage functional data analysis approach to examine global and local associations between along-tract diffusion metrics (FA, MD) and age, candidate genetic variants, and behavioural outcomes. First, global, tract-level associations were tested using penalised functional regression (PFR), modelling the scalar response, i.e., age, candidate genetic variants, or behavioural outcomes, against a functional term, i.e., along-tract diffusion metric pattern, while adjusting for sex and head motion (and ancestry PCs for genetic analyses). Multiple comparisons across the seven callosal tracts were controlled using Benjamini–Hochberg FDR correction for age- and behavioural-related PFR analyses (applied across the seven tracts for each behavioural factor score). For genetic analyses multiple comparisons were corrected using LD-aware significance thresholds.

For tracts showing significant global PFR associations, follow-up local, segment-specific associations were mapped using function-on-scalar regression (FOSR). For genetic analyses, FOSR was restricted to leading SNPs, i.e., independent variants that both passed the candidate gene-specific LD-aware significance threshold and survived clumping (r² < 0.1, 100 kb window). FOSR estimated smooth coefficient functions across the 98 tract segments while accounting for within-subject correlation. Segment-wise effect sizes (functional Cohen’s d) were extracted to quantify the magnitude of local effects. Significance was assessed at the segment level using the Benjamini–Bogomolov procedure.^83^

#### Comparison of along-tract with tract-average models

To explore the added value of a functional data analysis approach over linear regression (LM) models using simple tract average, we compared PFR and LM models using the Akaike Information Criterion (AIC), Akaike weights and related evidence ratios ^84^ (Supplementary Methods).

### Data availability

Data used in this study were obtained from the Philadelphia Neurodevelopmental Cohort (PNC), a collaboration between the Brain Behavior Laboratory at the University of Pennsylvania and the Center for Applied Genomics at the Children’s Hospital of Philadelphia (CHOP). PNC data are publicly available through the NIMH Database of Genotypes and Phenotypes (dbGaP; accession number phs000607). All softwares used in this study are openly available. Diffusion image preprocessing pipelines are available through QSIprep https://qsiprep.readthedocs.io/en/latest/ and TractSeg https://github.com/MIC-DKFZ/TractSeg/. Genetic data processing was completed using PLINK2 https://www.cog-genomics.org/plink/2.0/ and bcftools https://github.com/samtools/bcftools. The code for statistical analyses is openly available at https://gitlab.com/translationalml/cc_whitematter_developmental_cohort.

## Results

### Spatial modelling of callosal tract development

Based on AIC comparisons (Supplementary Tables S4 and S5), the best-fitting PFR models revealed significant global associations between along-tract FA patterns and age across all seven callosal tracts, and between along-tract MD patterns and age in all tracts except the splenium, with explained variance varying widely across tracts (Table 1).

For FA, model effect sizes (Cohen’s *f²*) ranged from small to medium across tracts, with values spanning 0.01 to 0.23. Notably, the strongest age associations were observed in the anterior midbody (*f²* = 0.23), genu (*f²* = 0.15), rostral body (*f²* = 0.15) and isthmus (*f²* = 0.13), whereas smaller yet significant effects were found in the rostrum, splenium and posterior mid-body (*f²* = 0.01–0.05). For MD, model effect sizes ranged from Cohen’s f² = 0.25 to 0.44 with large effect size observed in all callosal tracts, expect for the splenium showing only small and non-significant association (*f²* = 0.0008).

FOSR analyses revealed heterogeneous local age effects along each callosal tract (Figure 2, Supplementary Tables S6 and S7). Regarding FA (Figure 2, A), the rostrum displayed relatively stable, positive age associations with small variation across segments. In contrast, the genu, rostral body, anterior midbody, and isthmus showed a clear spatial gradient in age-related effects, with the strongest positive effect, i.e., increase FA with age, consistently emerging toward the distal ends of the tracts. The posterior midbody and splenium also showed more stable pattern of association along the tracts, with small effect size overall. With respect to MD (Figure 2, B), the patterns of age association were broadly similar across the rostrum, genu, rostral body, anterior midbody, and isthmus, with the largest and consistently negative effects emerging toward the distal portions of the tracts. Splenium’s along-tract MD pattern was not significantly associated with age.

**Figure 2.**
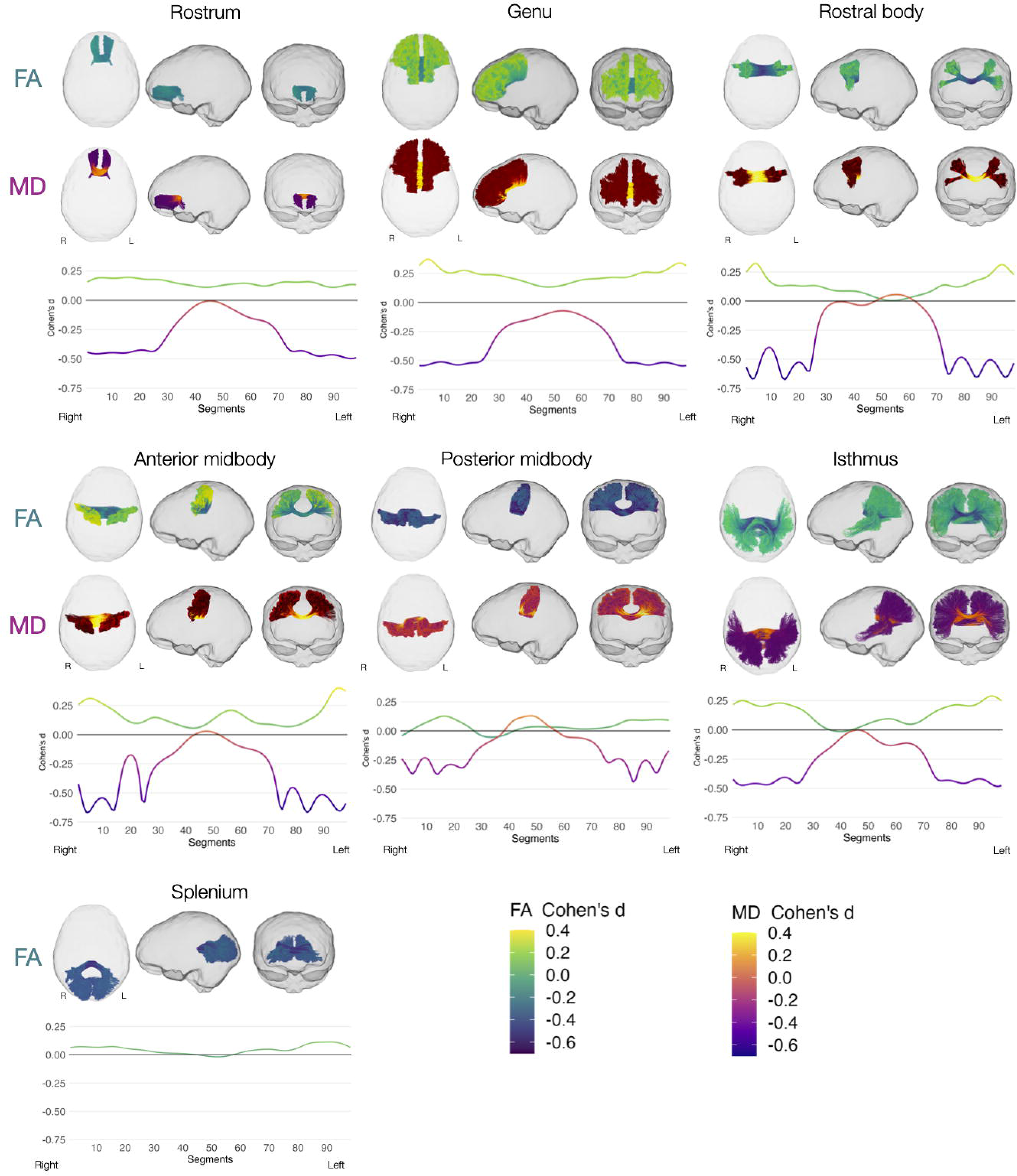
Age effects along callosal tract FA and MD metrics using function-on-scalar regression (FOSR) for the significant penalised functional regression (PFR) models. For each callosal tract (rostrum, genu, rostral body, anterior midbody, posterior midbody, isthmus, splenium), the glass-brain visualisations and corresponding line plots display the Cohen’s d estimated at each of the 98 callosal tract segments that quantify the magnitude of the along-tract effect of age on the given diffusion metric. Glass-brain visualisation: Each callosal tract is shown in an exemplar participant, with tracts coloured according to the local value of d according to FA and MD metrics. Axial, sagittal, and coronal views illustrate the spatial distribution of age effects along the tract. Line plots: The curve displays Cohen’s *d* across the 98 segments according to FA and MD. A horizontal reference line at *d* = 0 is included to facilitate interpretation of positive versus negative age effects. Line colour reflects the magnitude of d using the same colour scale as the glass-brain visualisations, enabling direct correspondence between the two representations for FA and MD.

PFR and tract-averaged linear models comparison using evidence ratios consistently favoured PFR models, with support ranging from weak to very strong across callosal tracts (Supplementary Table S8 and S9). The only exception was the splenium MD model, which showed no significant age association under either framework. In line with this, Cohen’s *f²* values were systematically higher for PFR models, or equivalent in the case of splenium MD.

### Along-tract association with candidate genetic variants

We examined along-tract associations for a total of 4809 genetic variants, including 1566 SNPs in ROBO1, 1251 SNPs in IQCJ-SCHIP1, 575 SNPs in NRP1 and 1417 SNPs in DCC.

Based on AIC model comparisons (Supplementary Tables S10 to S17), the best-fitting PFR models revealed significant associations between genetic variants in candidate genes and along-callosal metrics (*P* < gene-specific LD-aware threshold), with subsequent clumping showing independent lead SNPs predominantly in the rostrum (adjusted R² = 0.15–0.25, P = 2.2e-16–7.1e-05), isthmus (adjusted R² = 0.06–0.18, P = 2.2e-16–1.4e-04), and splenium (adjusted R² = 0.05–0.31, P = 2.2e-16–5.7e-05), (Supplementary Results, Supplementary Tables S18 to S26). Significant SNPs within each gene were largely tract-specific, with no overlap across callosal tracts. Functional annotation of independent lead variants was performed using intro for the subset of lead SNPs with the highest *adjusted R²* within each candidate gene across three predominant callosal tracts segments, i.e., rostrum, isthmus and splenium (Figure 3). Intronic variants were identified within DCC (rs62083512 [chr18:49922375], rs12953825 [chr18:49930196], rs1943098 [chr18:50404377]), ROBO1 [chr3:78913331], and IQCJ-SCHIP1 (rs114300450 [chr3:159046387]). Several variants were intergenic, falling within the ±100 kb: two for ROBO1 (rs572785555 [chr3:78549915] and rs181170008 [chr3:78565472]) and one for IQCJ-SCHIP1 (rs558636916 [chr3:158855955]). For NRP1, rs2131530723 [chr10:33466359:T/C] was annotated as a 3′ downstream variant located approximately 61 bp past the NRP1 transcription end site, while rs78043763 [chr10:33416046:T/C] was intergenic, approximately 50 kb upstream of the gene body.

**Figure 3.**
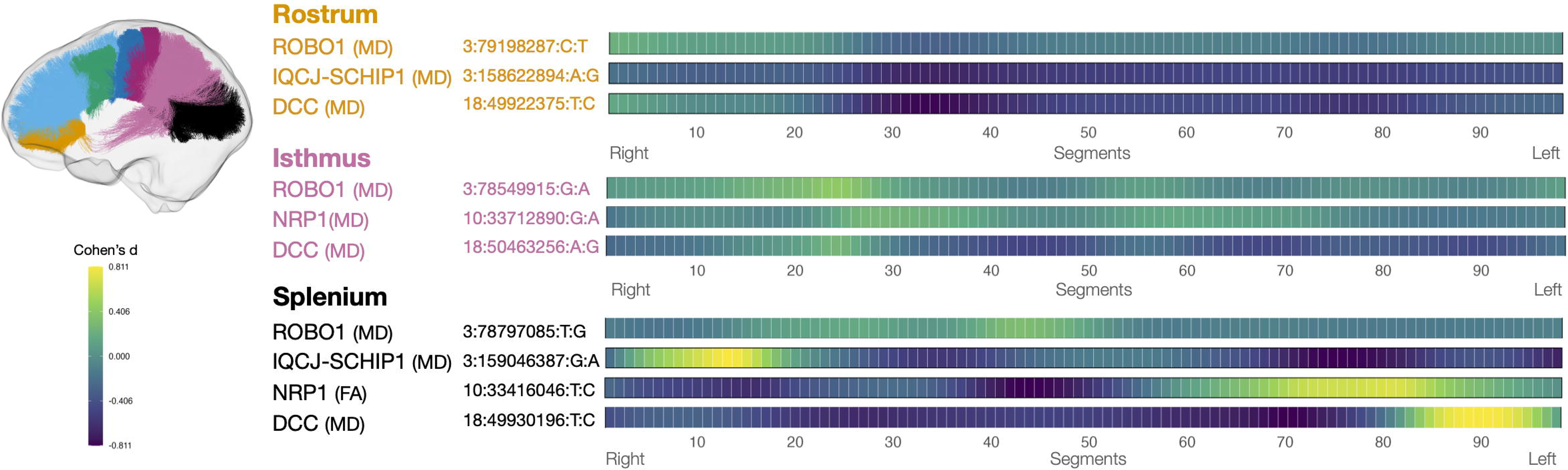
Significant associations between candidate gene variants and along-tract diffusion metrics. Global penalised functional regression (PFR) models identified significant associations between candidate gene variants and along-tract diffusion metrics (p < gene- specific LD-aware threshold), with independent leading SNPs after LD-based clumping predominantly for the rostrum, isthmus, and splenium. For each callosal tract, heatmap illustrates the function-on-scalar regression (FOSR)-derived local, segment-level patterns for each leading SNP selected as the variant explaining the greatest variance in the global PFR model. Rows correspond to individual leading SNPs, columns represent the 98 along-tract segments from right to left hemispheric endpoints, and colour intensity reflects the local effect size (Cohen’s *d*). FA: fractional anisotropy; MD: mean diffusivity.

FOSR analyses revealed that lead significant variants within selected genes were associated with spatially heterogeneous modulation of callosal microstructure, primarily within the rostrum, isthmus, and splenium, with effects varying across genetic variants (Supplementary Figures S4 to S7 and Supplementary Tables S27 to S30). In the rostrum, associations were spatially homogeneous across all four candidate genes, with moderate and broadly distributed effects along the full tract. In contrast, the isthmus and splenium showed gene-specific lateralised patterns, with stronger associations concentrated in either the right-most or left- most segments.

PFR and tract-averaged linear model comparison using evidence ratios showed that, across diffusion metrics, callosal tracts, and candidate genes, most model comparisons favoured the tract-averaged linear model with weak support (Supplementary Tables S33 and Table S34). When very strong and strong evidence was observed, it exclusively favoured PFR models. For models with strong evidence for PFR, *adjusted R²* values were higher for PFR than for LM models.

### Along-tract association with behavioural outcomes

Based on AIC comparisons (Supplementary Tables S35 and S36), the best-fitting PFR models revealed significant global associations but small effect size between along-tract callosal FA patterns and all four behavioural factors, with FA showing stronger associations than MD overall (Supplementary Tables S37 and S38, Figure 4). For complex cognition, significant associations with spatial variation in FA were observed across all callosal tracts except the anterior midbody (all FDR-corrected *P* ≤ 0.001; Cohen’s *f²* = 0.03–0.05). Memory and executive functioning were significantly associated with spatial variation in FA along all seven callosal tracts (all FDR-corrected *P* ≤ 0.01; *f²* = 0.007–0.05). For social cognition, significant associations with along-tract FA were observed specifically in the rostral body (FDR-corrected p = 0.04; *f²* = 0.006) and the posterior midbody (FDR-corrected *P* = 0.04; *f²* = 0.01). Regarding MD, significant associations for complex cognition with spatial variations in the genu, rostral body, anterior and posterior body, and isthmus were observed (FDR-corrected *P* ≤ 0.001 - 0.014; *f²* = 0.009–0.03). Executive functioning was significantly associated with spatial variation in the genu, anterior and posterior body (all FDR-corrected *P* ≤ 0.001; *f²* = 0.01–0.02). Finally, spatial variation in MD of the genu was associated with memory (FDR-corrected *P* = 0.028; *f²* = 0.01). It is worth noting that models incorporating along-tract spatial variation consistently explained more variance than those using averaged FA or MD values.

**Figure 4.**
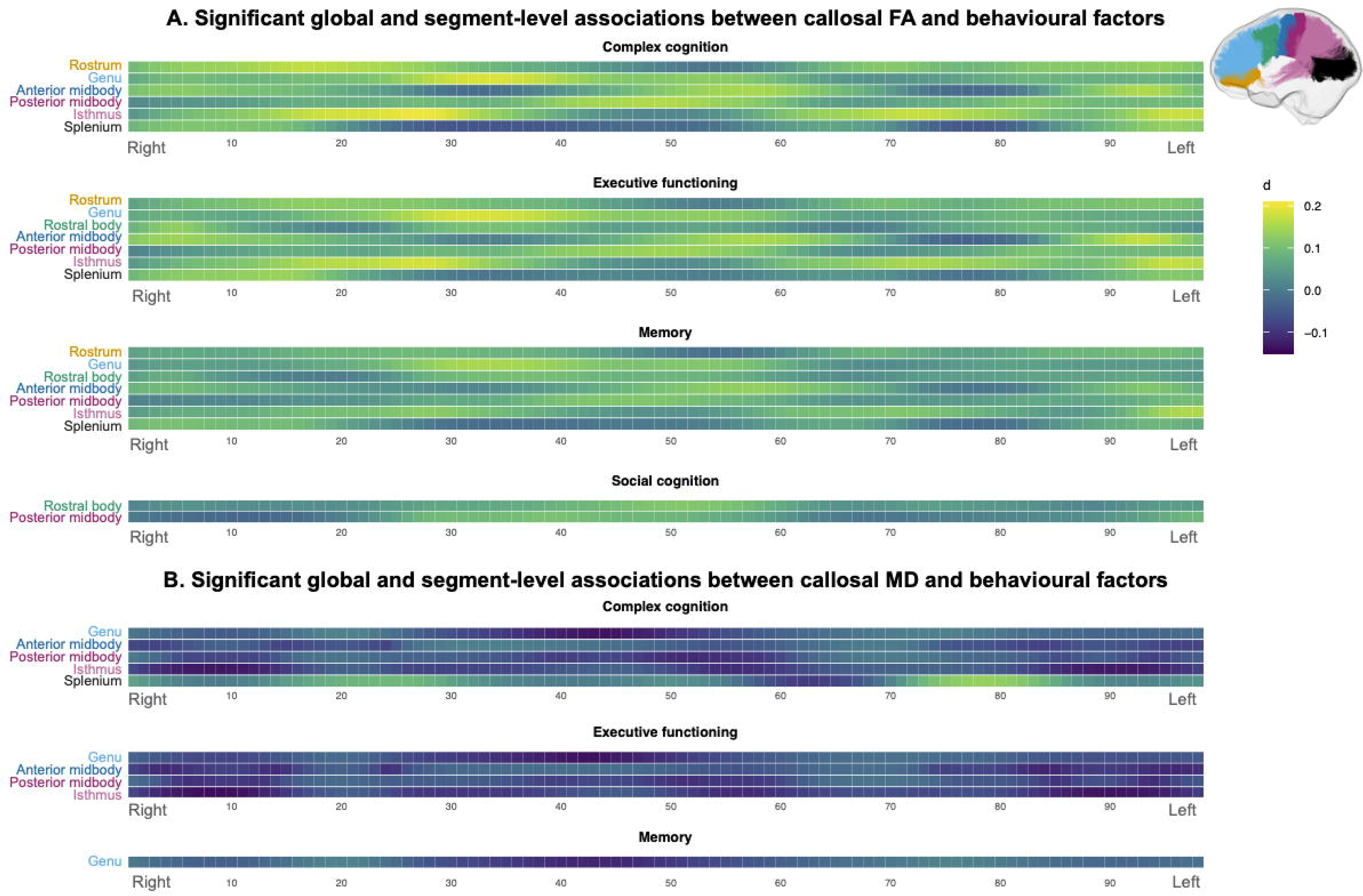
Significant global and segment-level associations between callosal FA/MD and behavioural factors. For each significant global association identified using penalised functional regression (PFR) between along-tract diffusion metrics and cognitive domains, heatmaps illustrate local segment-specific effects derived from function-on-scalar regression (FOSR). Heatmaps represent Cohen’s *d* estimates indicating the strength and direction of the association between behavioural performance and FA (A) or MD (B) across 98 tract segments (x-axis) within each significant callosal tract. Warmer colours indicate stronger positive effect sizes, whereas cooler colours indicate weaker or negative effects.

FOSR analyses revealed segment-level heterogeneity in the association between callosal FA and MD with behavioural factors (Supplementary Tables S39 and S40). Complex cognition and executive functioning showed the most heterogeneous patterns with pronounced variations in effect size along the callosal tracts, with both symmetric and asymmetric pattern of variations along tracts. Memory and social cognition showed a smoother and more homogeneous pattern, with less pronounced spatial fluctuations of effect size along the tracts.

PFR and tract-average linear models comparison showed that, across diffusion metrics, callosal tracts, and cognitive domains, PFR models were generally favoured (Supplementary Tables S41 and S42). Evidence ratios ranged from weak to very strong, with the strongest support for PFR observed in anterior and midbody callosal segments, most consistently for FA. Cohen’s *f²* values were typically higher for PFR models.

## Discussion

Considering the importance of the corpus callosum in human brain organisation and acknowledging the limitations of tract-averaged diffusion measures, we conducted a fine-grained characterisation of spatial variation along callosal tracts and examined how these patterns relate to developmental, genetic, and behavioural processes. Leveraging multimodal data and a functional data analysis approach, our findings revealed distinct midline-to-cortical variations of age-related diffusion metrics change across callosal tracts. Furthermore, spatial variation in callosal microstructure was modulated by genetic variants involved in early callosal axon guidance and was related to individual differences in behavioural outcomes. The findings also confirm that the additional model complexity introduced by a functional data analysis approach, in comparison to tract-average linear modelling, is justified by superior explanatory and predictive performance, underscoring the importance of explicitly modelling along-tract spatial variability.

Consistent with well-established age-related changes in tract-averaged diffusion metrics across callosal subdivisions, our results showed on average increases in FA and decreases in MD with age along the tracts.^47,85^ However, moving beyond averaged measures, we demonstrate that callosal maturation during childhood and adolescence follows a finely differentiated spatial pattern with each callosal subdivision exhibiting a unique spatial pattern of age-related effects. Among callosal tracts connecting frontal areas, the rostrum - linking predominantly orbitofrontal cortices - showed the most spatially stable association with age. While evidence show marked changes in orbitofrontal white-matter microstructure during development,^86^ these short callosal projection fibres appear to have a homogeneous maturation pattern. By contrast, other interhemispheric tracts connecting frontal regions (genu, rostral body, anterior midbody) and parietal regions (isthmus) demonstrated pronounced spatial heterogeneity in age-related changes, with the largest FA increases and MD decreases emerging toward distal segments near cortical terminations. Interpreting FA and MD as reflecting a measure of myelination,^43,47^ this pattern aligns with converging cellular and neuroimaging evidence showing that cortical myelination follows a prolonged, spatially non-uniform developmental trajectory specifically in these heteromodal and unimodal cortical areas. Radiocarbon birth-dating of oligodendrocytes indicates that cortical myelination remains remarkably dynamic from adolescence through adulthood, with oligodendrocyte numbers increasing until they plateau in the fourth decade of life.^87^ Complementary in vivo MRI studies indicate that adolescence is characterised by increases in intracortical myelin, particularly in mid-to-deep cortical layers of heteromodal and unimodal association regions,^88^ which correspond closely to cortical territories connected by these callosal fibres. Callosal fibres connecting sensorimotor and occipital cortices (posterior midbody and splenium) showed weaker age associations along their length, a pattern consistent with the earlier maturation and relative stability of these functional systems.^47,85,89^ These tracts also lacked the pronounced distal-end FA increases and MD decreases observed in heteromodal pathways, aligning with evidence of less prominently cortical myelinisation in idiotypic cortices. Visually, age-related effects in these posterior callosal tracts seem predominantly asymmetrical, suggesting that left- and right-sided fibres may follow partly distinct maturational pathways. Overall, these findings underscore that typical callosal maturation is not uniform but instead shaped by spatially differentiated patterns that vary both across callosal subdivisions and along their length.

Beyond age-related maturation, our findings also suggest that callosal spatial microstructural organisation is shaped by genetic variation in axon-guidance genes involved in early CC formation, including ROBO1, IQCJ–SCHIP1, NRP1, and DCC. We observed that variants within the selected axon-guidance genes were consistently associated with diffusion metrics variation in the rostrum, isthmus, and splenium—the most anterior and posterior portions of the CC. The selected genes are implicated in early commissural axon guidance, which begins at approximately 11–12 weeks of gestation.^6,18^ During this period, cingulate and neocortical fibres start forming the anterior CC, while the posterior CC is also forming, closely linked to the formation of the hippocampal commissure. Rather than a front-to-back formation, axonal guidance of the CC arises from these two distinct regions, which start to fuse around 13–14 weeks of gestation.^90,91^ Our findings suggest that genetic mechanisms active during these early stages of callosal formation may leave enduring spatial signatures more specifically in these early forming anterior and posterior callosal regions microstructure. Moreover, examination of the most significant variants confirmed that their associations were primarily located within the rostrum, isthmus, and splenium. Inspection of these top hits further revealed that most correspond to rare polymorphisms located within intronic regions of the candidate genes. For example, intronic variants were identified within DCC (rs62083512 [chr18:49922375], ROBO1 [chr3:78913331], and IQCJ-SCHIP1 (rs560590331 [chr3:158807945]. These findings therefore suggest that regulatory genetic variation within axon-guidance genes may modulate gene expression during early commissural development, ultimately contributing to inter-individual differences in callosal microstructural organisation. Notably, the majority of these variants are relatively rare (minor allele frequency <1%), supporting the possibility that subtle regulatory variation within axon-guidance pathways contributes to inter-individual variability in callosal microstructural organisation.

Extending these global findings, local association analyses indicated that genetic effects on callosal microstructure were spatially heterogeneous within tracts. The findings suggest that axon-guidance variants may shape local fibre properties within a tract and differentially affect distinct fibre segments, which may vary in axon diameter, packing density, and myelin-related characteristics.^92–94^ Moreover, the observation of both left and right spatial asymmetry in several tracts suggests a potential interaction between genetic influences and hemispheric asymmetry and specialisation, warranting further investigation.^3,95,96^

Having characterised the developmental and genetic correlates of callosal along-tract metrics, we next evaluated whether these spatial patterns were behaviourally meaningful. Across all four cognitive domains, which are essential for everyday functioning and academic success and are frequently disrupted in neurodevelopmental disorders, spatial variation in callosal FA showed significant associations with behavioural performance. Along-tract MD exhibited more limited associations, predominantly localised to central callosal tracts (genu, body, and isthmus) and primarily related to complex cognition and executive functioning. Such widespread association across callosal microstructure is consistent with the distributed interhemispheric integration across large cortical territories required by these domains.^97,98^ For along-tract FA in particular, segment-wise analyses revealed pronounced local variability in associations with complex cognition and executive functioning. These localised FA– behaviour relationships may reflect differences in how interhemispheric communication is balanced across hemispheres, strongly influenced by genetic and age-related factors, but this interpretation warrants further investigation.

Functional and LM model comparisons using AIC-based evidence ratios and effect size metrics broadly supported PFR across developmental, genetic and behavioural analyses. Taken together, these findings confirm that spatial variation along callosal tracts carries meaningful variance that is not captured by a single mean value, and that explicitly modelling this variation translates into improvements in explanatory power.

Our results should be interpreted in light of limitations. First, although the diffusion tensor model has been widely used and is recognised to yield robust and interpretable findings, it also comes with well-documented limitations. In particular, although DTI is sensitive to microstructural variation, similar changes in DTI metrics may arise from different underlying microstructural processes, limiting the specificity of their interpretation.^99^ Advanced diffusion models, such as diffusion kurtosis imaging (DKI) or fixel-based morphometry, could offer more specific biological insights into myelination or axonal density. However, the acquisition protocol used in the present study did not support the application of these higher-order models. Second, although TractSeg provides state-of-the-art tractography, the corpus callosum tracts derived with this method predominantly reflect homotopic interhemispheric connections. Recent work suggests that the human corpus callosum also contains heterotopic projections ^100^ that may contribute to developmental vulnerability. These heterotopic fibres are not fully captured using current tract parcellation approaches and may therefore limit our ability to characterise more complex patterns of interhemispheric structural connectivity. Finally, although the PNC offers a large developmental sample, our analyses remain cross-sectional. Longitudinal data would allow to delineate individual developmental trajectories of callosal microstructure, studying potential sensitive periods, and their association with behavioural outcomes.

To conclude, our findings converge on the CC as a spatially differentiated structure in which early genetic modulators and developmental constraints shape region- and segment-specific microstructural architecture that is behaviourally relevant through childhood and adolescence. By moving beyond tract-averaged measures, this spatially resolved framework highlights callosal along-tract organisation as a plausible intermediate phenotype linked to genetic variation, neurodevelopmental processes and behavioural heterogeneity. Such approach may help refine neurobiological models of developmental vulnerability and guide future work aiming to link genetic risk, white-matter organisation, and cognitive phenotypes in children with neurodevelopmental disorders.

## Acknowledgements

This work was supported by the Swiss National Science Foundation under the Ambizione Grant PZ00P1_208969 (to VS). Support for the collection of the data for Philadelphia Neurodevelopment Cohort (PNC) was provided by grant RC2MH089983 awarded to Raquel Gur and RC2MH089924 awarded to Hakon Hakonarson. Subjects were recruited and genotyped through the Center for Applied Genomics (CAG) at The Children’s Hospital in Philadelphia (CHOP). Phenotypic data collection occurred at the CAG/CHOP and at the Brain Behavior Laboratory, University of Pennsylvania.

## Supplementary material

Supplementary material is available at *Brain* online.

