## Supplementary Materials for "Mapping corpus callosum architecture: developmental, genetic, and cognitive correlates in youth"

### Supplementary Methods

#### dMRI Acquisition and processing

dMRI scans were acquired at University of Pennsylvania on a 3-T Siemens TIM Trio scanner (Siemens Medical Solutions), acquired using a twice-refocused spin-echo single-shot echo-planar imaging sequence sequence (FoV 240 × 240 mm; matrix 128 × 128 × 70; 64 gradient directions; b = 1000 s/mm$^2$; voxel size 1.875 × 1.875 × 2 mm) [1].

dMRI data preprocessing and reconstruction were performed using QSIPrep 0.19.1 [2], which is based on Nipype 1.8.6 ( [3, 4] RRID:SCR_002502).

Much of the text in the following two paragraphs was provided by *QSIPrep* under a CC0 license, and we made minor changes for succinctness. A total of 2 diffusion-weighted imaging (DWI) series in the j- distortion group were concatenated, with preprocessing operations performed on individual DWI series before concatenation. Any images with a b-value less than 100 s/mm$^2$ were treated as a b=0 image. MP-PCA denoising as implemented in MRtrix3's *dwidenoise* [5] was applied with a 5-voxel window. After MP-PCA, the mean intensity of the DWI series was adjusted so all the mean intensity of the b=0 images matched across each separate DWI scanning sequence. B1 field inhomogeneity was corrected using *dwibiascorrect* from MRtrix3 with the N4 algorithm [6] after corrected images were resampled. FSL (version 6.0.5.1)'s eddy was used for head motion correction and Eddy current correction [7]. Eddy was configured with a q-space smoothing factor of 10, a total of 5 iterations, and 1000 voxels used to estimate hyperparameters. A linear first level model and a linear second level model were used to characterize Eddy current-related spatial distortion. q-space coordinates were forcefully assigned to shells. Eddy was configured to separate field offset from subject motion. Shells were aligned post-eddy. Eddy's outlier replacement was run. Data were grouped by slice, only including values from slices determined to contain at least 250 intracerebral voxels. Groups deviating by more than 4 standard deviations from the prediction had their data replaced with imputed values. Final interpolation was performed using the jac method. FD using the implementation in Nipype (following the definitions by [8]) was calculated based on the pre-processed DWI and used as a measure of head motion. The DWI time-series were resampled to ACPC, generating a preprocessed DWI run in ACPC space with 1.875mm isotropic voxels.

The DTI model was applied to the resulting pre-processed DWI, and whole-brain maps of FA and MD were calculated for each participant. FA (between 0 and 1) is a measure of the directionality of diffusion that characterise the variance of the three eigenvalues pairs that represent the direction and magnitude of diffusivity along the three orthogonal axes ($v1$, $\lambda 1$; $v2$, $\lambda 2$; $v3$, $\lambda 3$). MD is the mean of the 3 eigenvalues and represents the average magnitude of diffusion [9]. Multi-tissue fibre response functions were estimated using the dhollander algorithm. Fiber orientation densities (FODs) were estimated via constrained spherical deconvolution (CSD, [10, 11]) using an unsupervised multi-tissue method [12, 13]. A single-shell-optimized multi-tissue CSD was performed (*mrtrix_singleshell_ss3t_noACT*) using MRtrix3Tissue (https://3Tissue.github.io), a fork of MRtrix3 [14]. FODs were intensity-normalized using *mtnormalize* [15]. Voxelwise fibre orientation density (FOD) peaks were then used as inputs for tract segmentation (described in the next section), which reconstructs streamlines, i.e., virtual trajectories approximating the course of anatomical white-matter bundles.

#### Tract segmentation

White matter bundles of the corpus callosum were segmented using *TractSeg* [16] for seven callosal subdivision including the Rostrum (denoted CC 1), Genu (CC 2), Rostral Body (CC 3), Anterior Midbody (CC 4), Posterior Midbody (CC 5) and Isthmus (CC 6) and the Splenium CC (CC 7). *TractSeg* is an automated toolbox using an encoder-decoder fully connected convolutional neural network (CNN) that processes voxelwise FOD peaks and then generates tract probability maps; these maps are then used to reconstruct tract-specific tractograms, i.e., sets of streamlines that approximate the spatial extent and orientation of the underlying white-matter bundle. The first three principal FOD peaks were extracted from the FODs image with MRtrix3 [14] and used as inputs into *TractSeg* to perform bundle segmentation, segmentation of the start and endpoints of the tracts, and tract orientation maps. Callosal tract segmentations were visually inspected and segment-wise tractometry (described below) of FA and MD for each callosal tract were examined to identify potential segmentation artefacts.

#### Tractometry

Each CC white-matter tract was divided into 100 segments using the assignment mapping approach described by Chandio et al. (2020) [17] and implemented into *TractSeg*. Every point of the tract was assigned to the closest points of the centroid on the model tract to create assignment maps/segments. Assignments are created in a common space (model tract), which ensures that the segment index corresponds to the same centroid across all individuals. A model tract is represented with 100 centroids uniformly distributed along the length of each tract in a common reference space (for more detailed procedure, see [17]). Importantly, this method uses all original points of the tracts, avoiding artificial resampling or reshaping. Tract segmentations were then applied to the diffusion metrics of interest, i.e., FA and MD maps, using the *Tractometry* implementation in *TractSeg* to segment-wise measures of callosal white-matter microstructure. This yielded 98 along-tract FA and MD values per tract (first and last segments are automatically removed due to unreliable estimates at tract endpoints).

**Genetic data processing and analysis**

*Genotype data acquisition*

Genotype data were obtained from dbGaP (phs000607.v2.p2). Biological samples (i.e., blood sample) from PNC participants who also completed the MRI were genotyped by the Center for Applied Genomics at the Children's Hospital of Philadelphia. Genotype were completed across participant in 8 batches using 7 different types of Affymetrix and Illumina arrays (i.e., Axiom, and Illumina HumanHap550 (v1, v3), *Human1M*-Duo (v3), Human610-Quad (v1), Genome-Wide Human SNP Array 6.0 and HumanOmniExpress) [18]. Supplementary Table S2 provide sample and SNP counts by genotyping array before and after QC, imputation and merge of genotype data.

*Quality control and cleaning of genotype data*

The genotype dataset was processed by array batch. Following previous studies using genetic data from the PNC dataset [19, 20], PLINK v2.0.0-a.6.13LM [21] was used to perform quality control, including the removal of individuals with >5% missing genotypes, single-nucleotide polymorphisms (SNPs) with >5% missingness, minor allele frequency <1%, or Hardy–Weinberg equilibrium p-value <1e-6. We standardised allele code across datasets by converting numeric allele encodings (1/2) to nucleotide format (A/C/G/T) using array-specific strand files from the manufacturer. Genome build harmonisation was performed by identifying datasets in NCBI36/hg18 (Omni, Axiom, Axiom_set2, Affy60) and converting them to GRCh37/hg19 with the UCSC *liftOver* tool (hg18ToHg19.over.chain.gz, accessed June 2025) [22]. Strand alignment was performed using *bcftools* version 1.21 against the 1000 Genomes Project Phase 3 reference panel [23], flipping mismatched SNPs and excluding ambiguous or multiallelic variants to ensure consistency across datasets. Allele normalisation was performed with *bcftools* using the hs37d5 reference genome (GRCh37) to ensure consistent allele representation across datasets [24]. Haplotype phasing was performed with *SHAPEIT5* (version 5.1.1) using GRCh37 genetic maps and chromosome-specific 1000 Genomes Phase 3 reference haplotypes.

*Imputation and merge of genotype data*

Phased genotypes were imputed using *IMPUTE5* (v1.2.0) with the 1000 Genomes Phase 3 reference panel and a b37 recombination map. Imputation was carried out separately for each dataset and each chromosome.

Imputed genotype data from multiple genotyping platforms were merged using *bcftools*. Only variants with high imputation quality (INFO score > 0.8) were retained prior to merging. Input VCF files, already normalized and annotated with allele counts, were sorted by genomic coordinates using *bcftools sort* and indexed with *bcftools index*. All datasets were subsequently merged into a single multi-array VCF using *bcftools merge.*

An unrelated sample was constructed and used for all subsequent analyses: for each family, the participant with the highest genotyping completeness across the four candidate gene regions was retained.

*Genetic Ancestry and population structure correction*

To account for population stratification [25], genetic ancestry was assessed using principal component analysis (PCA) on the merged imputed genotype dataset. The merged VCF was converted to PLINK format, duplicate variants were removed, and unique variant identifiers were assigned. Linkage disequilibrium (LD) pruning was performed using a sliding window approach ($r^2 < 0.2$). PCA was conducted using PLINK2 v2.0.0-a.6.13LM on LD-pruned variants and the first 20 principal components were computed. Based on the examination of the variance explained by each PC (Supplementary Figure S2) and previous studies [20], the first ten ancestry PCs were included as covariates in subsequent statistical models.

*Candidate gene coordinates*

As described in the introduction section, four genes implicated in callosal axonal guidance were selected for the current candidate gene association study: NRP1, DCC, ROBO1 and IQCJ-SCHIP1. Gene coordinates (chromosome, start, end) were obtained from the Ensembl GTF annotation file

Homo_sapiens.GRCh37.87.gtf.gz (https://ftp.ensembl.org/pub/grch37/current/ gtf/homo_sapiens/). For each candidate gene, the annotated gene interval was retrieved and

candidate genetic variants were defined as SNPs located within the gene body and extended by ±100 kb flanking regions of ROBO1, IQCJ-SCHIP1, NRP1 and DCC.

*LD structure and gene-based significance thresholds*

SNPs with zero variance and, where applicable, fewer than 100 complete genotype–phenotype observations were excluded. For each candidate gene, LD matrices were computed as Pearson correlation coefficients (r) between SNP pairs using R (v4.4.3) (Supplementary Figure S3). To correct for multiple testing while accounting for LD correlation among SNPs, we applied an LD-aware Bonferroni correction per gene. Rather than assuming independence across all tested variants, the effective number of independent tests ($M_{eff}$) was derived from LD matrices using Gao's method, implemented in the *SimpleM* R package, and used to compute corrected significance thresholds [26–28] (Supplementary Table S3). To identify independent association signals within each candidate gene region, significant variants (p value < candidate gene-specific LD-aware threshold) were subsequently clumped using *snp_clumping* from the *bigsnpr* R package [82], with in-sample LD computed from study genotype data, an $r^2$ threshold of 0.1, and a 100 kb window.

**Statistical analyses**

All statistical analyses were conducted in R (v4.4.3).

All along-tract functional data analysis used a two-stage functional data analysis framework that models both global and local effects of spatial variation in callosal tract metrics. This approach allowed to examine associations with age, candidate genetic variants, and behavioural outcomes (post-hoc analyses).

*Along-tract developmental models*

We adopted a two-stage functional data analysis framework, first modelling global associations between along-tract pattern and age using penalized functional regression, and then mapping local specific age effects along the tract using function-on-scalar regression.

To model the global association between age and along-tract diffusivity metrics (FA and MD), we employed penalized functional regression (PFR), a scalar-on-function regression approach [29–31]. PFR enables the characterisation of associations between a scalar response (age) and a functional term (along-tracts metrics). PFR models were fitted using the *pfr* function from the *refund* package version 0.1.37 [32]. In this framework, the functional term is represented using spline basis functions, and the functional regression parameters (i.e., coefficient functions) are estimated as penalized splines to allow smooth yet flexible modelling of their effects [30]. In line with the implementation of PFR in the refund package and to ensure methodological consistency across functional models, tract patterns were presmoothed using $fpca.sc$ to obtain smoothed functional predictors before fitting the PFR model [30, 32, 33]. Seven separate PFR models were fitted, one for each CC tract, with along-tract FA as the functional term, age as a scalar term, and sex and mean FD (head motion) as covariates. The same procedure was applied to MD, resulting in seven additional models.

The penalised functional regression models were specified for each CC tract as:

$$Y_i = \gamma_0 + \gamma_1 \cdot sex_i + \gamma_2 \cdot motion_i + F\{X_{i(\cdot)}\} + \varepsilon_i$$

Where $Y_i$ is the scalar outcome age, $\gamma_0$ is the intercept, and $\gamma_1$ and $\gamma_2$ represent the regression coefficients for sex and head motion, respectively. The term $X_{i(\cdot)}$ denotes the tract pattern for participant $i$ across 98 along-tract segments. The term $F\{X_{i(\cdot)}\}$ is the functional term that captures the full along-tract profile by estimating a smooth coefficient function $\beta(s)$ across the 98 segments, allowing the association with age to vary continuously along the tract rather than being summarised by a single scalar coefficient. It is specified as a linear functional regression term, and estimated under a penalisation framework to control smoothness and avoid overfitting. Finally, $\varepsilon_i$ represents the residual error. Model fit was compared across different numbers of spline basis functions ($k = 20,30,40,50,60$) using the Akaike Information Criterion (AIC). Previous work suggest that the choice of k is not critical as long as it is sufficiently large to capture the maximum complexity of the regression function [30, 34]. For each tract, the model with the lowest AIC was selected as the optimal specification for subsequent analyses. Statistical inference focused on the functional term $F\{X_{i(\cdot)}\}$, which captures the association between the along-tract diffusion profile and age. The significance of this functional effect was assessed using the approximate F-test reported in the *pfr* model summary for the linear functional term. P-values were adjusted for multiple comparisons using the Benjamini–Hochberg procedure to control the false discovery rate (FDR) across the 7 callosal tracts.

For tracts showing a significant global association between the functional term and age in the PFR models (FDR-corrected p-value < .05), we then employed a function-on-scalar regression (FOSR) approach to identify tract segments where diffusion metrics vary as a function of age [35, 36]. FOSR treats the along-tract pattern as a functional response and estimates smooth coefficient functions describing the effect of a scalar predictor of interest (age) at each tract segment. FOSR models were fitted using the *fosr* function from the refund package version 0.1.37 in R version 4.4.3 (2025-02-28). To maintain methodological coherence with the PFR analyses, tract patterns were presmoothed using *fpca.sc*, ensuring identical preprocessing across PFR and FOSR models. The *fosr* function was then applied using the same spline basis dimension (k) that was selected for the corresponding PFR model, thereby matching basis parameters across global and local analyses.

For participant *i* at tract segment $s$, the FOSR models were specified as:

$$Y_{i(s)} = \beta_{0(s)} + \beta_{1(s)age_i} + \beta_{2(s)sex_i} + \beta_{3(s)motion_i} + u_i + \varepsilon_{i(s)}$$

where $Y_{i(s)}$ denotes the diffusion metric (FA or MD) for participant *i* at tract segment *s* (across 98 along-tract segments). $\beta_0$ represents the mean tract pattern (i.e. intercept), while $\beta_{1(s)}$, $\beta_{2(s)}$, $\beta_{3(s)}$ are smooth coefficient functions that describe how covariates age, sex, and head motion influence the diffusion signal at each tract segment. The subject-specific random intercept $u_i$ accounts for within-participant correlation across tract segments, and $\varepsilon_{i(s)}$ denotes the residual error at each segment *s*.

For each of the 98 callosal segments, functional Cohen's d at each segment location $s$ were extracted to quantify the magnitude of the along-tract effect of age on the diffusion metric, specified as:

$$\mathrm{d(s)} = \frac{[\,\beta_{\mathrm{age(s)}} \times \mathrm{SD(age)}]}{\mathrm{SD}(\varepsilon(\mathrm{s}))}$$

where $\beta_{\mathrm{age}(s)}$ is the estimated coefficient function from the FOSR model, $\mathrm{SD(age)}$ is the standard deviation of the age predictor, and $\mathrm{SD}(\varepsilon(\mathrm{s}))$ is the residual standard deviation of the functional model at location $s$.

Significance was assessed at the segment level using the Benjamini–Bogomolov procedure [85]: segment-wise p-values were adjusted using the Benjamini–Hochberg method and compared against a selection-adjusted threshold $\alpha_{L2} = \alpha \times \frac{R_1}{m_1}$, where $\alpha_{L2}$ is the significance threshold for segment-level (Level 2: FOSR) analyses, $\alpha$ is the global significance level (i.e., 0.05), $R_1$ is the number of significant tract-level (Level 1: PFR) models, and $m_1$ is the total number of tract-level models tested.

*Along-tract association with candidate genetics variants*

To investigate whether specific genetic variants were associated with local variation in callosal microstructure, the same two-stage functional data analysis framework used for age was applied.

First, global associations between along-tract diffusivity metrics (FA and MD) metrics and each genetic variant of interest were modelled using PFR. This scalar-on-function approach tests associations between the genetic variant (coded additively as 0/1/2 copies of the minor allele) and the along-tracts diffusion metrics functional term. Subsequent PFR modelling procedures mirrored those used in the age analyses. In short, tract patterns were presmoothed using *fpca.sc* to obtain smoothed functional predictors before fitting the PFR model [30, 32, 33]. Seven separate PFR models were fitted, one for each CC tract, with along-tract FA as the functional term, genetic variant as a scalar term, and the following covariates: sex, mean framewise displacement (head motion), and the first ten ancestry PCs to account for population structure. The same procedure was applied to MD. The penalised functional regression models were specified for each CC tract as:

$$Y_i = \gamma_0 + \gamma_1 \cdot sex_i + \gamma_2 \cdot motion_i + \gamma_3 \cdot PC1_i + \gamma_4 \cdot PC2_i + \gamma_5 \cdot PC3_i + \gamma_6 \cdot PC4 + \gamma_7 \cdot PC5_i + \gamma_8 \cdot PC6_i + \gamma_9 \cdot PC7_i + \gamma_{10} \cdot PC8_i + \gamma_{11} \cdot PC9_i + \gamma_{12} \cdot PC10_i + F\{ X_{i(\cdot)}\} + \varepsilon_i,$$

Where $Y_i$ is the scalar outcome genetic variants, $\gamma_0$ is the intercept, and $\gamma_1$ , $\gamma_2$ and $\gamma_3$ to $\gamma_{12}$ represent the regression coefficients for sex, head motion, and the ten ancestry PCs, respectively. The term $X_{i(\cdot)}$ denotes the tract pattern for participant $i$ across 98 along-tract segments, and $F\{ X_{i(\cdot)}\}$ is the functional term that captures the tract pattern. It is specified as a linear functional regression term and estimated under a penalisation framework to control smoothness and avoid overfitting. Finally, $\varepsilon_i$ represents the residual error. Model fit was compared across different numbers of spline basis functions ($k = 20,30,40,50,60$) using the Akaike Information Criterion (AIC), as described in the previous section. The association between genetic variant for each candidate gene and along-tract diffusion metrics were considered statistically significant if their p-values exceeded the gene-specific LD-aware significance threshold.

For tracts showing a significant global association between the functional term and genetic variant in the PFR models (based on LD-aware significance threshold), we then employed a function-on-scalar regression (FOSR) approach to identify tract segments where diffusion metrics vary as a function of age [35, 36]. Subsequent FOSR modelling steps followed the same procedures as in the age analyses with the genetic variant entered as the scalar predictor, including functional pre-smoothing of tract metrics, fitting FOSR models with sex, head motion and the ten ancestry PCs as covariates, applying the same spline basis dimension selected in the corresponding PFR model (best AIC criteria), extracting segment-wise Cohen's d. To avoid rank-deficiency in the FOSR models, ancestry PCs exhibiting extremely low variance (standard deviation < $1\times10^{-3}$) were excluded prior to model fitting. Significance was assessed at the segment level using the Benjamini–Bogomolov procedure [85]: segment-wise p-values were adjusted using the Benjamini–Hochberg method and compared against a selection-adjusted threshold $\alpha_{L2} = \alpha \times \frac{R_1}{m_1}$, where $\alpha_{L2}$ is the significance threshold for segment-level (Level 2: FOSR) analyses, $\alpha$ is the global significance level (i.e., 0.05), $R_1$ is the number of significant tract-level (Level 1: PFR) models, and $m_1$ is the total number of tract-level models tested.

*Along-tract association with behavioural outcomes*

Finally, to examine whether behavioural outcomes were associated with variation in callosal microstructure, we applied the same two-stage functional data analysis framework used for age and genetic variants. First, global associations between each behavioural measure and along-tract diffusion metrics (FA and MD) were modelled using PFR, treating the behavioural outcome as the scalar predictor. All modelling steps—including functional pre-smoothing of tract metrics, specification of tract-wise PFR models with sex and motion as covariates, optimisation of spline basis dimension via AIC, and FDR correction for each tract—followed the procedures described above. For significant PFR models, local segment-wise associations were then identified using FOSR, with the behavioural outcome entered as the scalar predictor. As in previous analyses, tract metrics were presmoothed, FOSR models were fitted using the same spline basis dimension selected in the corresponding PFR model and extracting segment-wise Cohen's d. Significance was assessed at the segment level using the Benjamini–Bogomolov procedure [85]: segment-wise p-values were adjusted using the Benjamini–Hochberg method and compared against a selection-adjusted threshold $\alpha_{L2} = \alpha \times \frac{R_1}{m_1}$, where $\alpha_{L2}$ is the significance threshold for segment-level (Level 2: FOSR) analyses, $\alpha$ is the global significance level (i.e., 0.05), $R_1$ is the number of significant tract-level (Level 1: PFR) models, and $m_1$ is the total number of tract-level models tested.

*Comparison of along-tract with tract-average models*

To explore the added value of a functional data analysis approach over linear regression (LM) models using simple tract average, we compared PFR and LM models using the Akaike Information Criterion (AIC) and Akaike weights [86]. For each tract and outcome, both models were fitted using the same sample and covariate set, with the sole difference being whether callosal diffusion entered as a 98-segments functional pattern (PFR) or a scalar average (LM: single mean value across the 98 segments). Akaike weights and evidence ratios were computed following Burnham & Anderson (2002):

ΔAIC is defined as the difference in AIC between the two candidate models (PFR and LM):

$$\Delta AIC = AIC_{LM} - AIC_{PFR}$$

Positive ΔAIC indicates better fit for PFR and negative ΔAIC indicates better fit for LM. Akaike weights (w) for the PFR model were computed as:

$$w_{PFR} = \frac{exp(-0.5 \cdot \Delta AIC)}{1 + exp(-0.5 \cdot \Delta AIC)}$$

Akaike weights for the LM model were computed as:

$$w_{LM} = 1 - w_{PFR}$$

Where $w_{PFR}$ and $w_{LM}$ represent the probability that a given model is the best-supported candidate given the data. The evidence ratio (ER) was then computed as:

$$ER = \frac{w_{PFR}}{w_{LM}}$$

ER quantify how much more likely one model is to be the best-supported candidate relative to the other. When ER > 1, evidence favoured PFR; when ER < 1, evidence favoured LM. Strength of evidence was classified following Burnham & Anderson (2002), with ER (or 1/ER when ER < 1): between 1 and 2.7, ≥ 2.7, ≥ 10, and ≥ 100 corresponding to weak, substantial, strong, and very strong support for the favoured model, respectively.

### Supplementary Figures

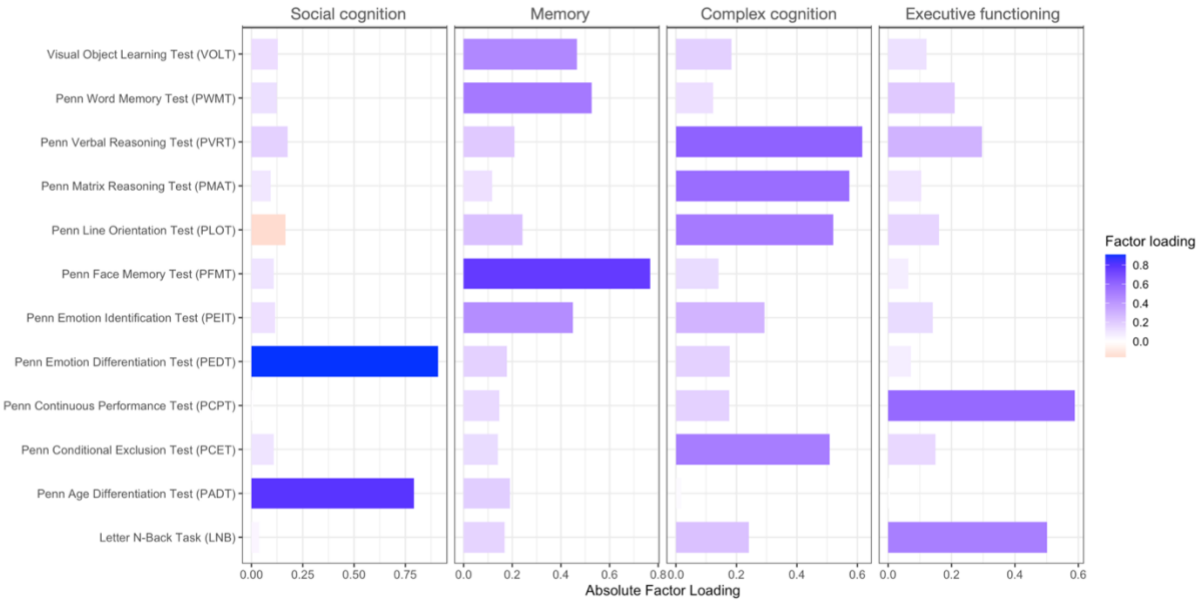


**Figure S1**. Factor structure of neurocognitive efficiency measures derived from exploratory factor analysis. Bar plots show the absolute factor loadings of the 12 cognitive efficiency measures on the four latent factors identified by exploratory factor analysis: executive functioning, episodic memory, complex cognition, and social cognition. Colour intensity reflects the magnitude of factor loadings, with darker shades indicating stronger associations between individual tests and their corresponding latent domain.

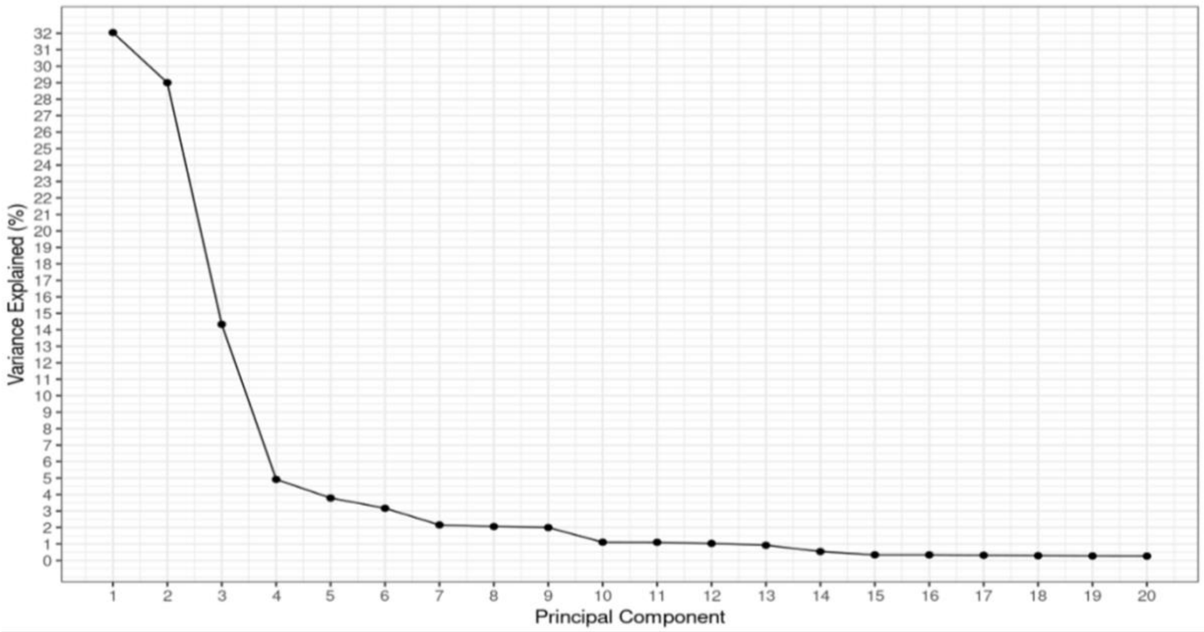


**Figure S2 –** Scree plot of principal component analysis (PCA) for genetic ancestry estimation. The screeplot shows the proportion of variance explained by the first 20 principal components derived from PCA on linkage disequilibrium (LD)–pruned, merged imputed genotype data. Based on the variance explained and consistency with previous studies [20], the first ten ancestry principal components were retained as covariates to control for population stratification in subsequent statistical analyses.

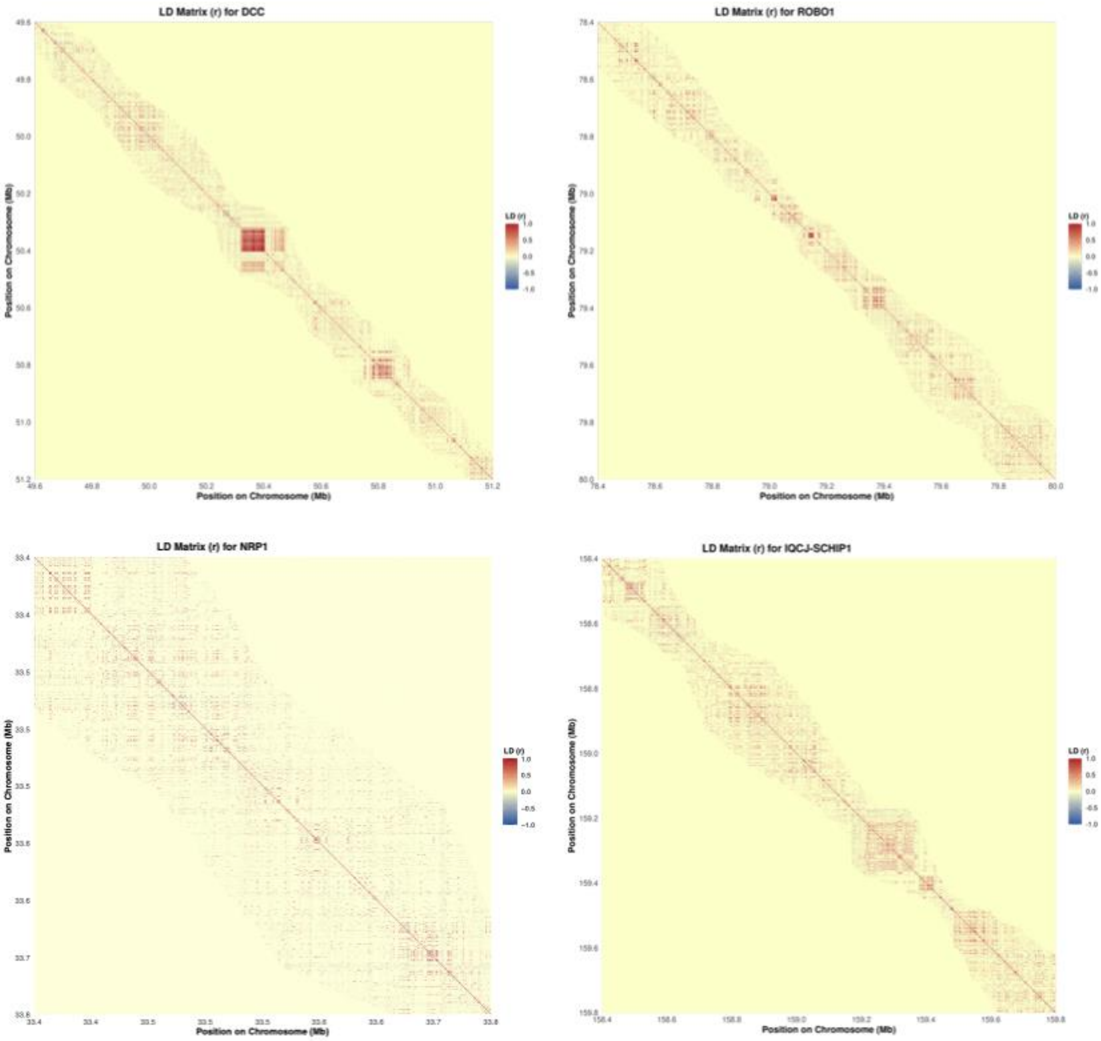


**Figure S3 -** Linkage disequilibrium (LD) structure within candidate gene regions. Heatmaps show pairwise linkage disequilibrium (Pearson correlation coefficient, r) between SNPs within the extended genomic regions (gene body ±100 kb) of the four candidate genes DCC, NRP1, IQCJ-SCHIP1, NRP1 and ROBO1. SNPs are ordered by genomic position, with colour intensity reflecting the magnitude and direction of LD (red = positive correlation, blue = negative correlation).

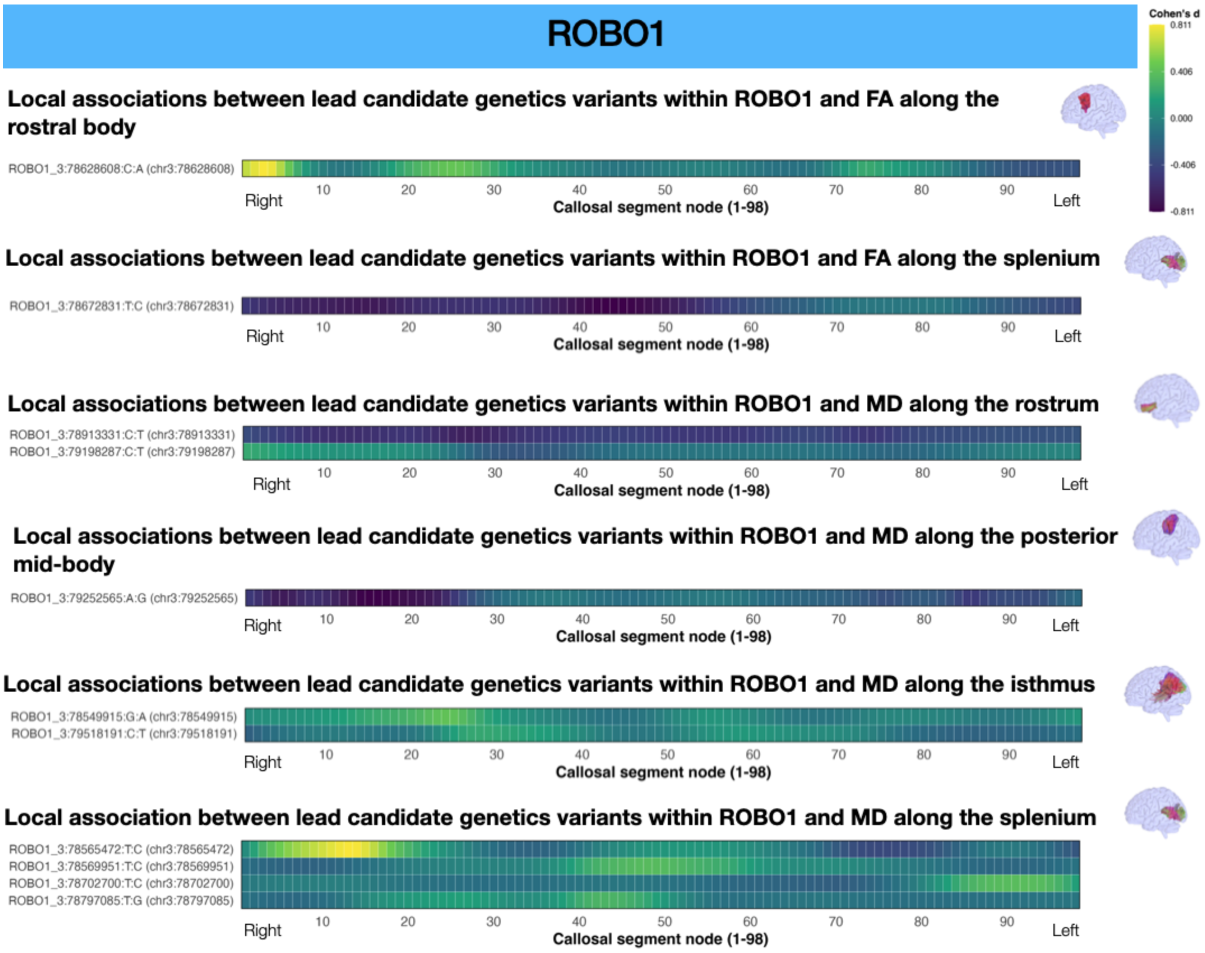


**Figure S4.** Local associations of independent leading genetic variants based on PFR — first thresholded at p < candidate gene-specific LD-aware threshold, then identified through LD-based clumping — within ROBO1 with callosal tract FA and MD metrics using function-on-scalar regression (FOSR). The curve displays Cohen's *d* estimated at each of the 98 callosal tract segments that quantify the magnitude of the along-tract effect of genetic variants within ROBO1 along the diffusion metric. A horizontal reference line at d = 0 is included to facilitate interpretation of positive versus negative age effects. Line colour reflects the magnitude of d.

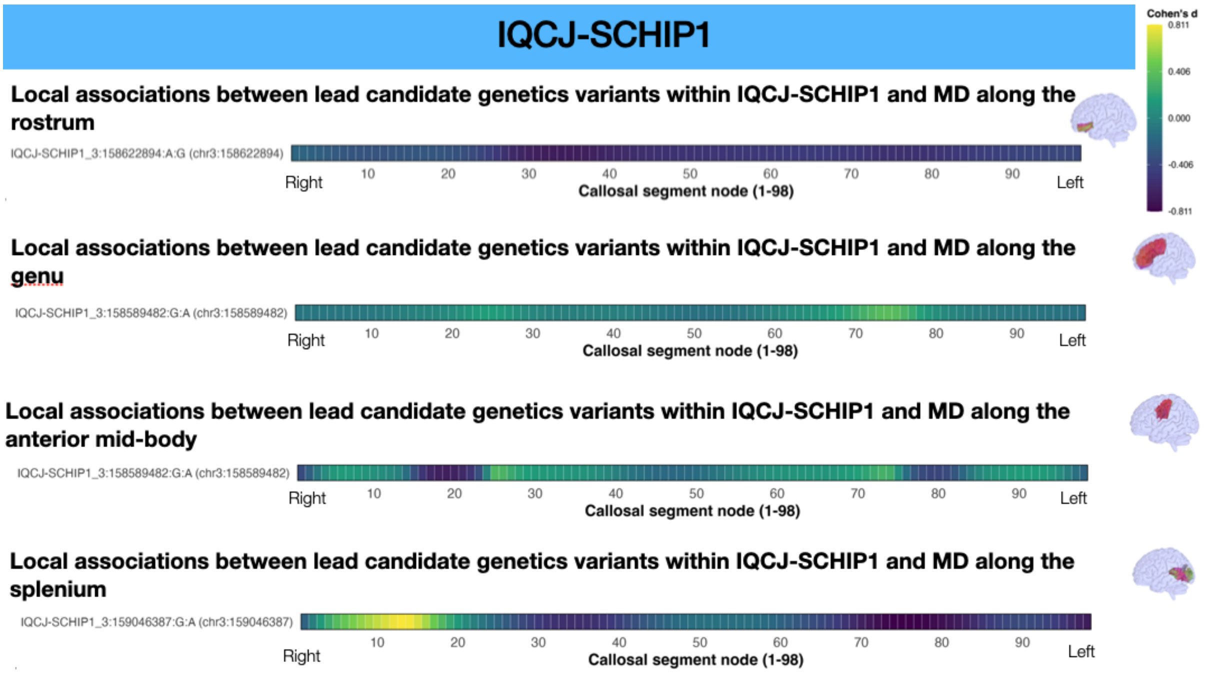


**Figure S5.** Local associations of independent leading genetic variants based on PFR — first thresholded at p < candidate gene-specific LD-aware threshold, then identified through LD-based clumping — within IQCJ-SCHIP1 with callosal tract FA and MD metrics using function-on-scalar regression (FOSR). The curve displays Cohen's *d* estimated at each of the 98 callosal tract segments that quantify the magnitude of the along-tract effect of genetic variants within ROBO1 along the diffusion metric. A horizontal reference line at d = 0 is included to facilitate interpretation of positive versus negative age effects. Line colour reflects the magnitude of d.

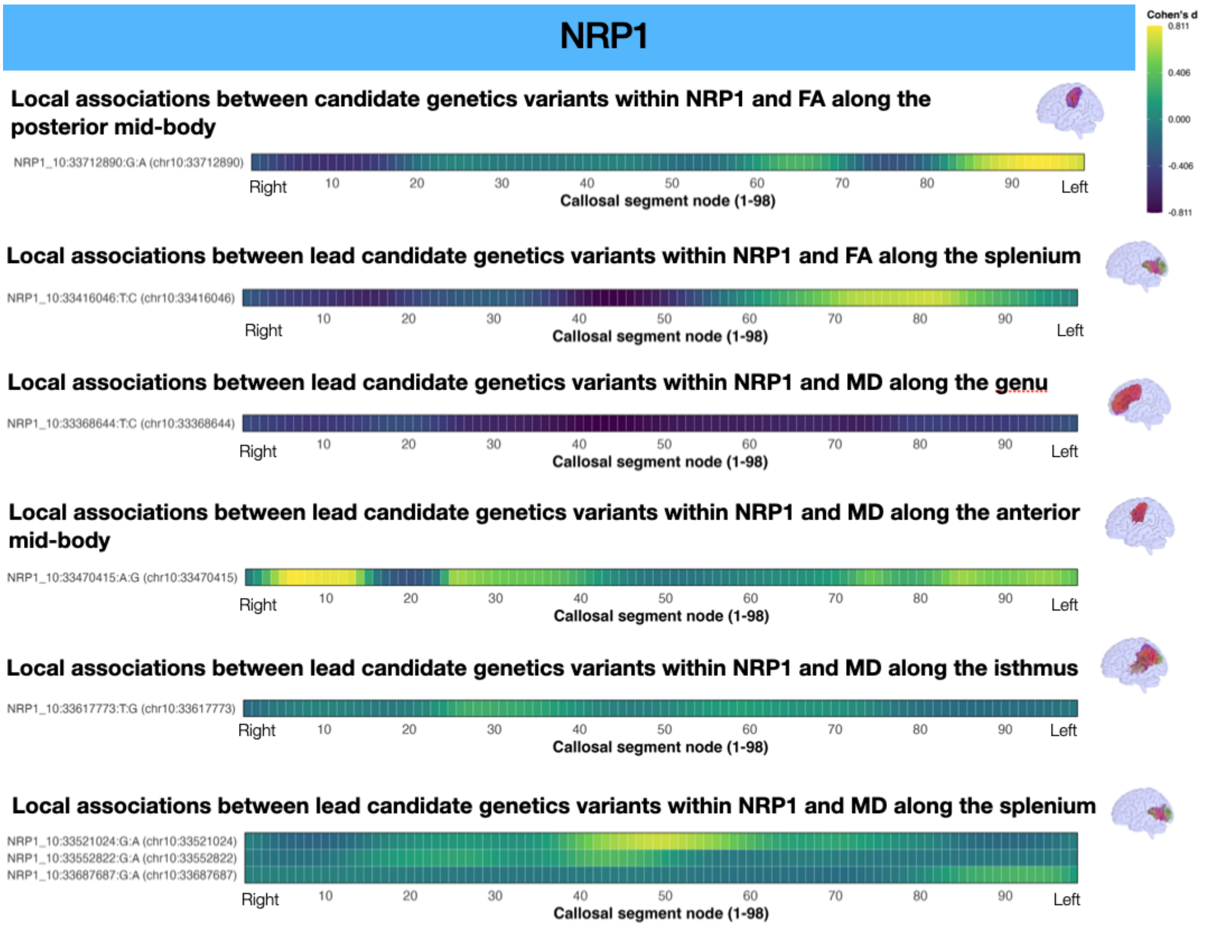


**Figure S6.** Local associations of independent leading genetic variants based on PFR — first thresholded at p < candidate gene-specific LD-aware threshold, then identified through LD-based clumping — within NRP1 with callosal tract FA and MD metrics using function-on-scalar regression (FOSR). The curve displays Cohen's *d* estimated at each of the 98 callosal tract segments that quantify the magnitude of the along-tract effect of genetic variants within ROBO1 along the diffusion metric. A horizontal reference line at d = 0 is included to facilitate interpretation of positive versus negative age effects. Line colour reflects the magnitude of d.

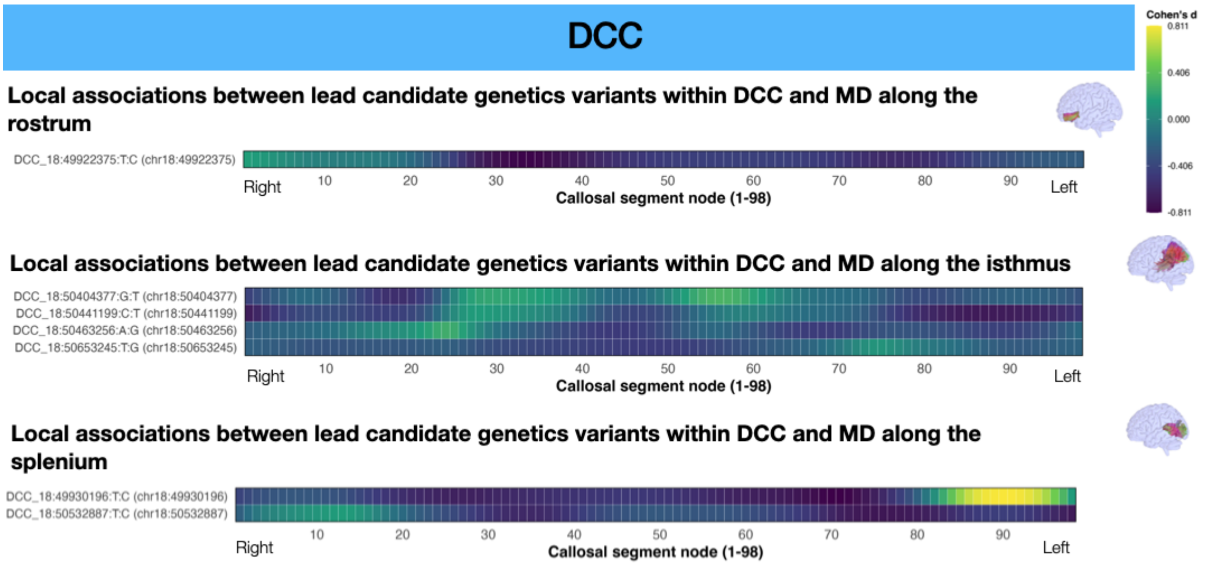


**Figure S7.** Local associations of independent leading genetic variants based on PFR — first thresholded at p < candidate gene-specific LD-aware threshold, then identified through LD-based clumping — within DCC with callosal tract FA and MD metrics using function-on-scalar regression (FOSR). The curve displays Cohen's *d* estimated at each of the 98 callosal tract segments that quantify the magnitude of the along-tract effect of genetic variants within ROBO1 along the diffusion metric. A horizontal reference line at d = 0 is included to facilitate interpretation of positive versus negative age effects. Line colour reflects the magnitude of d.
